# Hepatocyte-derived extracellular vesicles regulate liver regeneration after partial hepatectomy

**DOI:** 10.1101/2024.06.25.600679

**Authors:** Mina McGinn, Christopher Rabender, Ross Mikkelsen, Vasily Yakovlev

## Abstract

While significant progress has been made in understanding different aspects of liver regeneration, the molecular mechanisms responsible for the initiation and termination of cell proliferation in the liver after massive loss or injury of liver tissue remain unknown. The loss of liver mass affects tissue-specific mitogenic inhibitors in the blood, which in turn regulate the proliferation of remaining hepatocytes and liver regeneration. Although well described in a number of publications, which inhibitory substances or “sensor molecules” control the regeneration mechanisms to properly maintain liver size remain unknown. Extracellular vesicles (EVs) are nano-sized, membrane-limited structures secreted by cells into the extracellular space. Their proposed role is stable intercellular carriers of proteins and RNAs, mostly micro-RNA, from secreted to recipient cells. Taken up by the recipient cells, EVs can significantly modulate their biological functions. In the present study, using *in vivo* and *in vitro* models, we demonstrate that hepatocyte proliferation and liver regeneration are regulated by EVs secreted by hepatocytes into the bloodstream. This regulation is carried out through a negative feedback mechanism, which explains the very precise regeneration of liver tissue after massive damage. We also demonstrate that an essential component of this mechanism is RNA carried by hepatocyte-derived EVs. These findings open up a new and unexplored area of biology regarding the mechanisms involved in the homeostasis regulation of various constantly renewing tissues by maintaining the optimal size and correct ratio between differentiating and proliferating cells.

## Introduction

How constantly renewing tissues maintain an optimal size and a delicate balance between differentiating and proliferating cells is an intriguing question which remains unanswered. Some 80 years ago a series of now classic studies were published which all suggested that within any cell line the functionally active differentiated cells may signal their existence to the mitotically active precursor cells by means of a tissue or a cell-line specific inhibitory substance^1-4^. Application of this negative feedback concept to the autoregulation of cell proliferation and tissue regeneration was very appealing. Later the principal characteristics of these inhibitory substances were postulated ^5-11^: a) they inhibit mitosis both in vitro and in vivo; b) their action is reversible and they are not cytotoxic; c) they are synthesized by mature cells of the tissue upon which they act, released from cells and circulate in the blood stream and in humoral fluids; d) they are tissue-specific, but species-unspecific. One of the most known and frequently used model to study the regulation of cell proliferation is a rodent partial hepatectomy (PHx). The liver is known for its remarkable ability to precisely regenerate after massive tissue loss or injury^12^. Regeneration of the liver is studied by performing the PHx to remove 2/3 of the liver mass in rodents (rats and mice)^13-15^. The entire process of liver regeneration following PHx takes up to 10-14 days in rodents during which the remnant liver lobes grow through extensive proliferation of all hepatic cell types. Following regeneration, the liver grows back precisely to its original mass and does not exceed it^16^. Moreover, in liver transplant experiments in which, after PHx, the liver was transplanted from small animals to large ones and *vice versa* (for example, from baboons to humans, from small dogs to large dogs), the liver after transplantation was always restored in proportion to the size of the recipient animal^17^. Other studies have shown that loss of liver mass affects tissue-specific mitogenic inhibitors in the blood, which in turn regulate the proliferation of remaining hepatocytes and liver regeneration ^18-20^. Although well described in numerous publications, what inhibitory substances or “sensor molecules” control the regeneration mechanisms to properly maintain liver size remain unknown.

Extracellular vesicles (EVs) are nanosized, membrane-limited structures secreted by cells into the extracellular space^21,22^. EVs provide a new mode of cell-to-cell communication, in which their cargo is transferred from a donor to a recipient cell, leading to changes in gene expression and cellular function^23-26^. Micro-RNAs (miRNAs) are small (22–24 nucleotides) non-coding regulatory RNAs that regulate gene expression at the post-transcriptional level and are the main regulators of virtually all cellular processes. MiRNAs bind to complementary sequences in the 3′-untranslated region (3′UTR) of target mRNAs, leading to either translational repression or target degradation of the specific mRNA ^27^. One of the most recent findings is that miRNAs exist in EVs and these EVs’ miRNAs can be functionally delivered to target cells^28^. In the present study, using *in vivo* and *in vitro* models, we demonstrate that hepatocyte proliferation and liver regeneration are regulated by EVs secreted by hepatocytes into the bloodstream. This regulation is carried out through a negative feedback mechanism, which explains the very precise regeneration of liver tissue after its massive damage. An essential component of this mechanism is RNA carried by hepatocyte-derived EVs (HD-EVs).

## Results

To assess proliferative activity in the mouse liver after surgery, we applied the most commonly used biomarkers of cell proliferation: Cyclin D1 and PCNA. Western blotting and immunohistochemistry were used to analyze the overall expression level and localization of biomarkers in liver tissue. We demonstrated that on the 2^nd^ day after PHx, the period of greatest proliferative activity occurs, in which almost all hepatocytes of the remaining liver lobes were synchronically involved (Figure 1a-c). By the 10^th^ day after the PHx, when the original liver volume was restored, the expression of proliferation biomarkers and the percentage of proliferating hepatocytes had returned to the level of the intact (Control) animals. Next, we examined how the concentration of HD-EVs in the blood changed over the period of liver regeneration after PHx. Asialoglycoprotein receptor 1 (ASGR1) protein is a well-established hepatocellular protein, and, as it was shown previously, can serve as a biomarker for the HD-EVs^29,30^. To determine the concentration of HD-EVs in the mouse plasma, we used a double-staining method (Figure 1c). DiB, a lipophilic cationic dye, was used to stain all EVs isolated from mouse plasma, which allowed us to distinguish true EVs from other particles similar in size to EVs. Labeling of EVs with Alexa Fluor 647 (AF647)-conjugated anti-mASGR1 antibodies allowed identification of HD-EVs. By double-staining method we demonstrated that changes in HD-EVs concentration in the blood were inversely proportional to the level of proliferation activity in liver tissue after PHx (Figures 1d,e).

**Figure 1.**
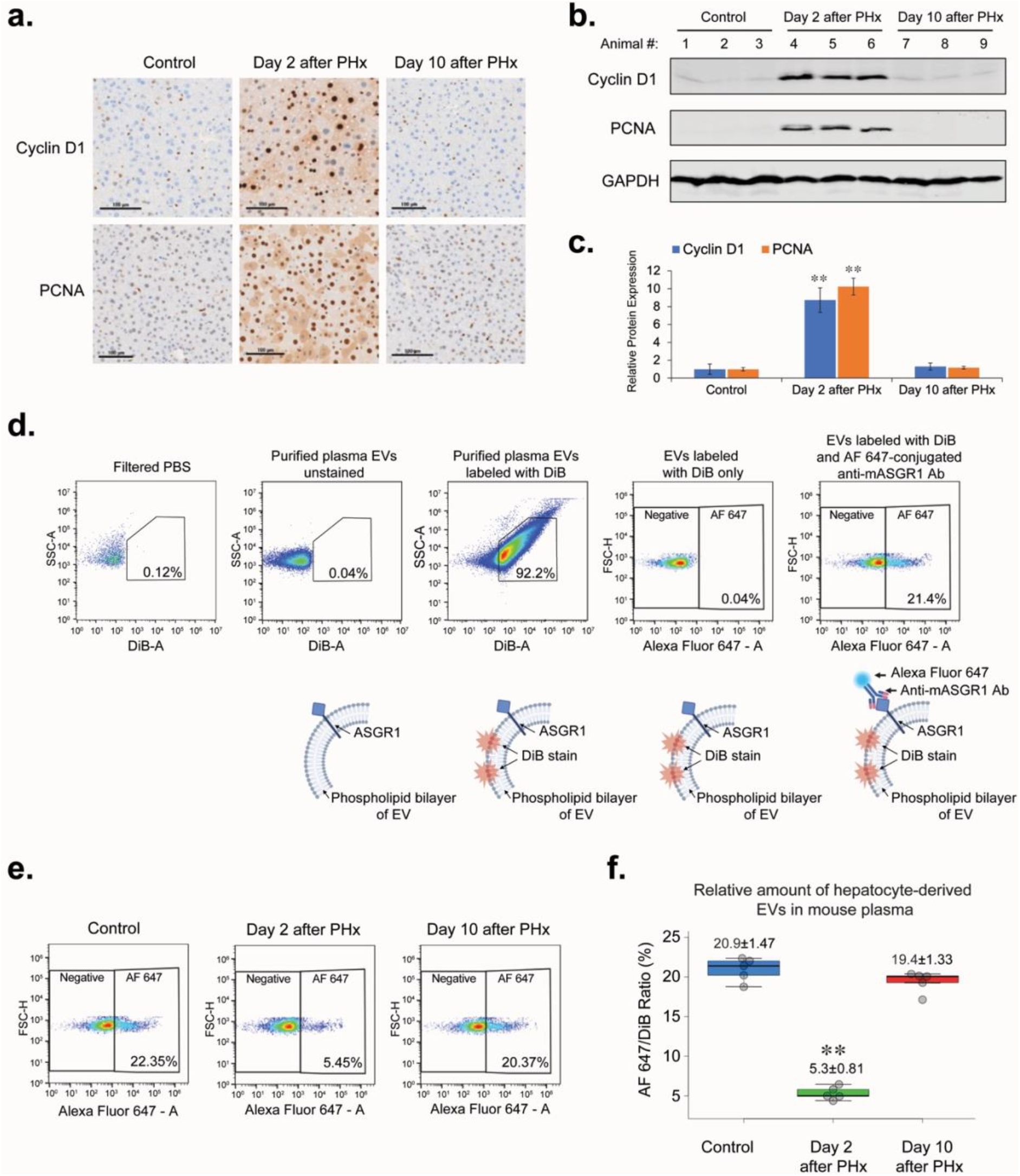
PHx affects liver hepatocyte proliferation and concentration of hepatocyte-derived EVs (HD-EVs) in mouse plasma. **a**. IHC for proliferation markers PCNA and Cyclin D1 in liver tissue of non-surgery control mice and at the different time-points after PHx. Scale bar = 100 µm. **b**. Western blot analysis of liver tissue lysates with Cyclin D1, PCNA, and GAPDH for different time-points after PHx and non-surgery control (n = 3 for each group). **c**. Graph with quantification of the Western blot analysis shown on Fig. 1b. ** - p-value < 0.001. **d**. Flow cytometry analysis of HD-EVs labeled with DiB and AF647-conjugated Anti-mASGR1 Ab and schematic representation of EVs labeling. **e**. Flow cytometry analysis of HD-EVs concentration in plasma of non-surgery control animals and in the different time-points after PHx. **f**. Graph with quantification of the flow cytometry analysis shown in Fig.1d. n = 6 animals in group. ** - p-value < 0.001.

In our next experiments we used EVs purified from commercially available C57BL/6 mouse plasma by the size exclusion chromatography (SEC). Ponceau S staining and nanoparticle tracking analysis of various fractions after extraction of EVs from mouse plasma by SEC showed that fractions #1-4, containing the bulk of EVs, were free from contamination by plasma proteins (Figure 2a,b). The same amount of total protein from the pooled fractions of EVs (fractions #1-4) and plasma proteins (fractions 8-11) were analyzed by Western blot (Figure 2c). The absence of specific EV markers (CD9, CD63, and TSG101) in the pooled protein fractions and lack of albumin (the most abandoned plasma protein) in the pooled EVs fractions demonstrates an efficient separation of EVs from plasma proteins. This method allowed recovery of >97% of purified EVs from mouse plasma. The average plasma concentration of EVs in intact healthy mice is 2-4×10^11^/ml (Figure 2e). In our experiments, in order to achieve an adequate response, when introducing purified EVs into the cell culture medium or into the blood of mice, we used concentrations close to the concentration of EVs in the plasma of the intact, healthy mice. Therefore, the purified exosomes were pre-concentrated before use (Figure 2e). Transmission Electron Microscopy analysis demonstrated that the concentration step didn’t change the size, shape, and integrity of EVs (Figure 2d).

**Figure 2.**
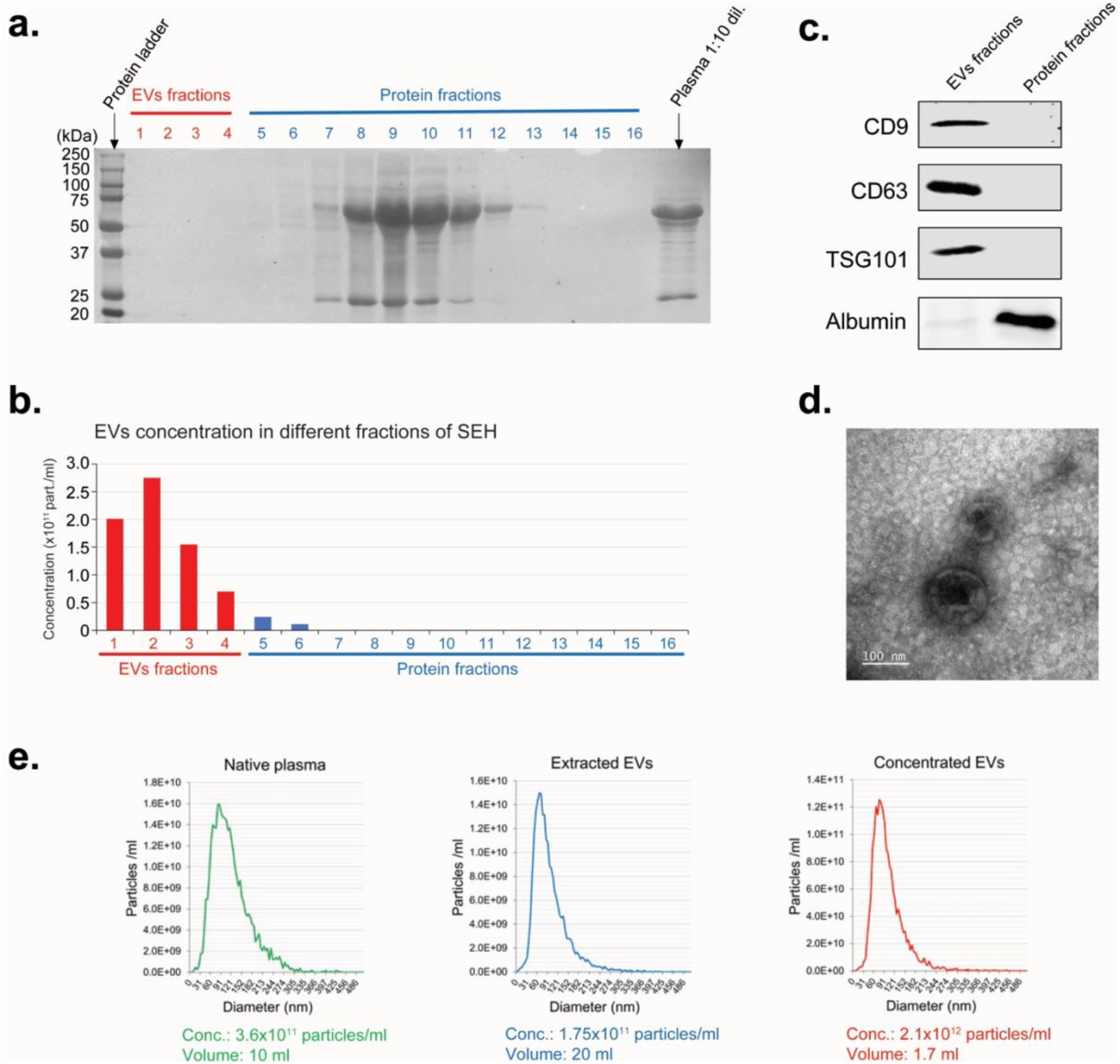
Extraction of EVs from mouse plasma. **a**. Ponceau S staining of the different plasma fractions after extraction of EVs from mouse plasma by SEC. **b**. Nanoparticle tracking analysis of EVs in different plasma fractions (as shown in Fig. 2a) after extraction of EVs by the SEC. **c**. Western blot analysis of EV-specific protein markers. Equal amount of total protein was loaded for combined EVs fractions (#1-4), and Protein fractions #8-11 (as shown in Fig. 2A). **d**. Electron microscopy analysis for the extracted EVs. Scale bar = 100 nm. **e**. Nanoparticle tracking analysis of EVs in the native plasma, after extraction from plasma by the SEH (Extracted EVs), and after concentration step (Concentrated EVs).

Our next experiments were focused on assessing the level of hepatocyte-derived and non-hepatocyte-derived EVs uptake by mouse hepatocytes (AML-12 cell line) was assessed *in vitro* (Figure 3a). We generated the HD-EV-depleted fraction of total plasma EVs (ASGR1-negEVs) by immunoprecipitation of HD-EVs with anti-ASGR1 Ab (Figure 3b). Incubation of normal mouse hepatocytes (AML-12) with DiL-stained non-hepatocyte (ASGR1-neg) EVs (NH-EVs) and global plasma EVs (GP-EVs) demonstrated that the uptake of GP-EVs, which contain HD-EVs and NH-EVs fractions, by AML-12 cells was significantly higher than that of NH-EVs only (Figure 3c). Thus, hepatocytes uptake HD-EVs more actively than EVs from other sources. Incubation of AML-12 cells with GP-EVs resulted in a dose-dependent inhibition of the cell cycle, with accumulation of cells in the G1 phase (Figures 3d,e), and reduction in expression of the proliferation biomarkers PNCA and Cyclin D1 (Figures 3f,g). It was previously demonstrated that Heparin blocks EVs’ cellular uptake. Our preliminary study also demonstrated a dose-dependent inhibition of GP-EVs’ uptake by AML-12 cells treated with heparin (Extended data, Figure S1). We determined that the optimal regimen for heparin is incubation at a dose of 5 µg/ml for 2 hours. This regimen significantly reduced GP-EVs uptake and did not affect the cell cycle of AML-12 cell line. Preincubation of GP-EVs and AML-12 cells with heparin prevented inhibition of proliferation of total plasma EVs (Figure 3d-g). Unlike GP-EVs, incubation of AML-12 cells with NH-EVs did not affect the cell cycle activity and expression of the proliferation biomarkers (Figure 3d-g). This data indicates that HD-EVs are responsible for the suppression of mouse hepatocyte proliferation *in vitro*.

**Figure 3.**
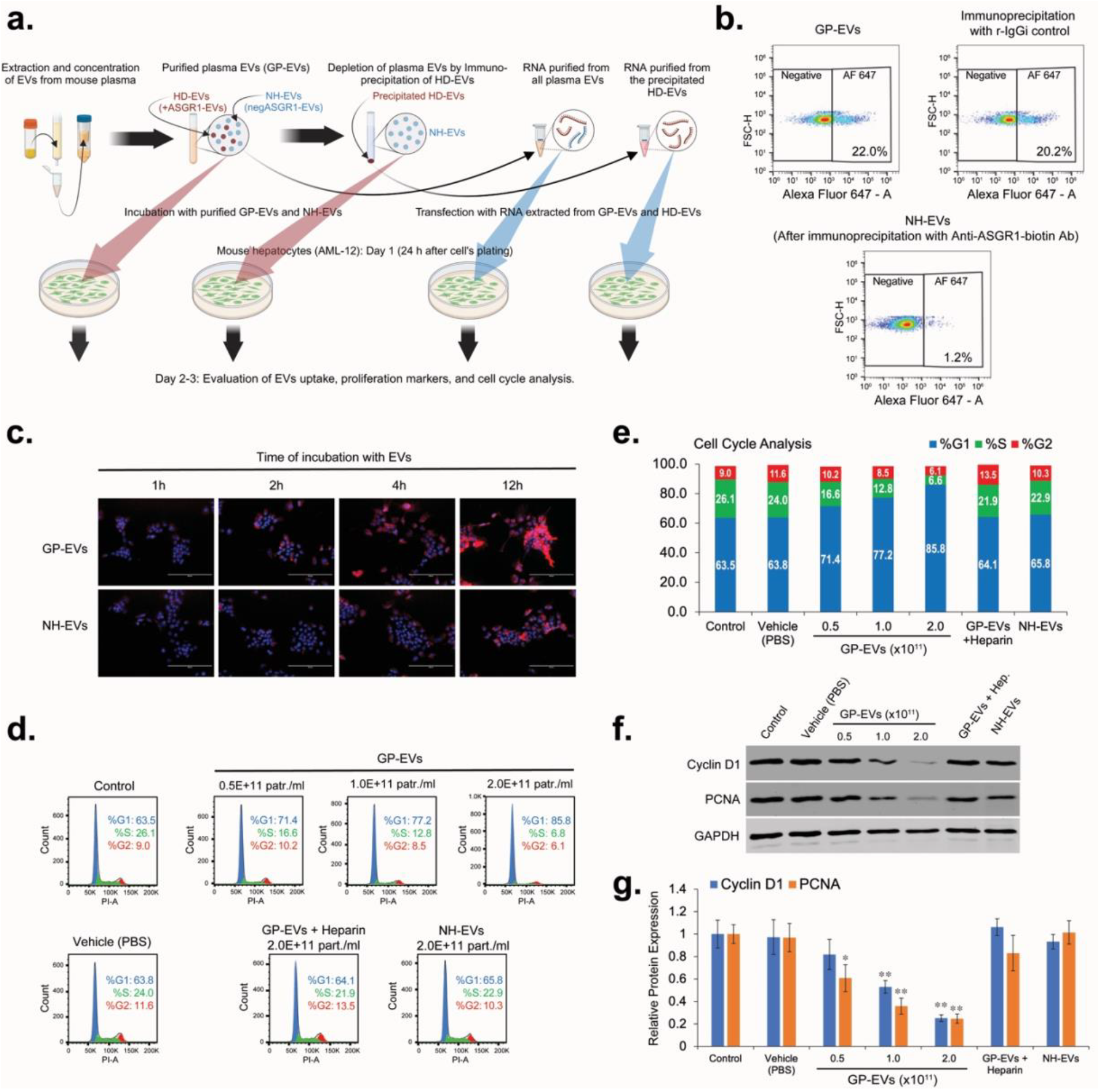
HD-EVs blocks proliferation of mouse hepatocytes (AML-12) *in vitro*. **a**. The schematic illustration of the in vitro experiments. **b**. flow cytometry analysis for EVs double-labeled with DiB/AF647-conjugated Anti-mASGR1 in the global plasma EVs (GP-EVs), after immunoprecipitation with biotinylated Anti-ASGR1 Ab (NH-EVs), and after immunoprecipitation with biotinylated rabbit IgG isotype (r-IgGi) control. **c**. EVs uptake assay for the GP-EVs and for the NH-EVs. 24 hours after seeding, AML-12 cells were incubated for different periods of time with two different types of DiL-labeled EVs at a final concentration of 2×10^11^ particles/ml. At the end of the incubation time, the cells were washed, fixed, and stained with mounting media with DAPI. Scale bar = 200 µm. **d**. Cell cycle analysis for AML-12 cells incubated with vehicle (PBS) and with different concentrations of GP-EVs extracted from mouse plasma. As negative controls: i) AML-12 cells and GP-EVs were preincubated for 2 hours with 5 µg/ml of Heparin; ii) AML-12 cells were incubated with NH-EVs. **e**. Graph representation of the cell cycle analysis shown in Fig. 3d. **f**. Western blot analysis for expression of Cyclin D1, PCNA, and GAPDH in lysates of AML-12 cells treated as shown in Fig. 3d. Shown is representative data from three independent experiments. **g**. Graph with quantification of the Western blot analysis shown in Fig. 3f. n = 3 independent experiments. * - p-value < 0.05; ** - p-value < 0.005.

To determine the role RNA carried by HD-EVs in the regulation of hepatocyte proliferation, total RNA (including miRNAs) was extracted from GP-EVs and HD-EVs immunoprecipitated with anti-ASGR1antibodies. AML-12 cells were transfected with RNA extracted from EVs 24 h after plating. 48 hours after transfection, cells were analyzed for cell cycle distribution and expression of proliferation biomarkers Cyclin D1 and PCNA. Transfection with RNA isolated from GP-EVs caused a dose-dependent arrest of AML-12 cells in the G1 phase (Figure 4a,b), which was accompanied by a significant decrease in the expression of Cyclin D1 and PCNA proteins (Figure 4c,d). Transfection with RNA isolated from HD-EVs demonstrated a more potent block of AML-12 proliferation compared to RNA from GP-EVs: the effect of 100 pM of HD-EVs-extracted RNA on AML-12 proliferation was more pronounced than effect of 500 pM of the total EVs-derived RNA (Figure 4a-d).

**Figure 4.**
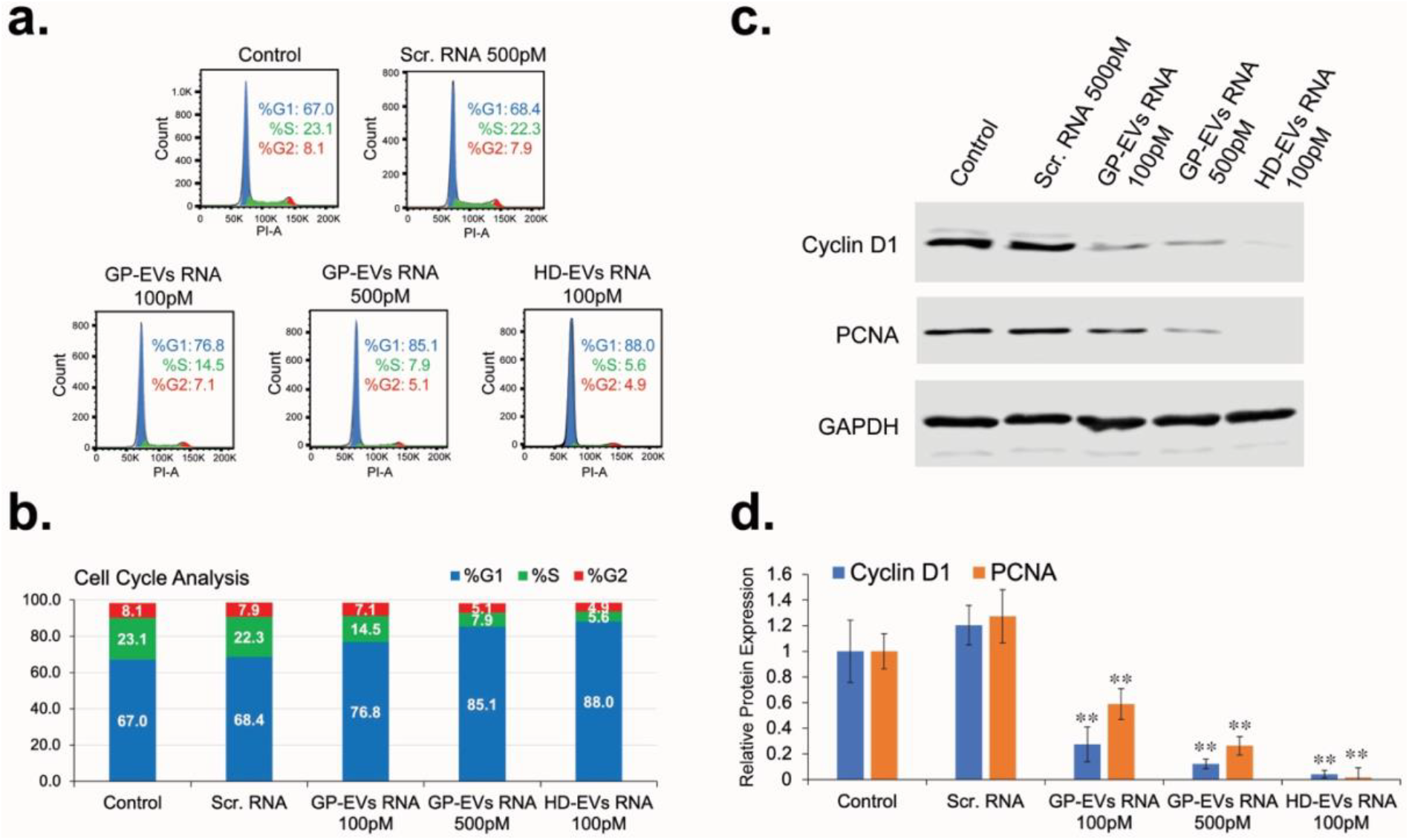
RNA extracted from HD-EVs blocks proliferation of mouse hepatocytes (AML-12) in vitro. **a**. Cell cycle analysis for AML-12 cells transfected with RNA extracted from GP-EVs, HD-EVs (ASGR1+ EVs), or with Negative Control siRNA. **b**. Graph representation of the cell cycle analysis shown in Fig. 4a. **c**. Western blot analysis for expression of Cyclin D1, PCNA, and GAPDH in lysates of AML-12 cells transfected as shown in Fig. 4a. Shown is representative data from three independent experiments. **d**. Graph with quantification of the Western blot analysis shown in Fig. 4c. n = 3 independent experiments. ** - p-value <0.005.

To evaluate the effect of EVs and RNA they carry on proliferation of hepatocytes *in vivo*, we used the two-thirds PHx mouse model (Figure 5a). DiR-stained GP-EVs can be detected in different organs of mice soon after intravenous (IV) injection, however, 24 hours later, a strong signal continues to be detected only in the liver (Figure 5b). After conducting PHx, animals received two sequential IV injections (12 hours and 24 hours after surgery) with: vehicle (PBS), GP-EVs, or NH-EVs. 36 hours after PHx animals were euthanized and liver samples were examined by IHC and Western blot analyses (Figure 5c-e). Injection with GP-EVs significantly attenuated PHx-stimulated increase of expression of Cyclin D1 and PCNA in a mouse liver. Unlike GP-EVs, IV injections with NH-EVs showed almost no impact on expression of Cyclin D1 and PCNA in liver tissue after PHx.

**Figure 5.**
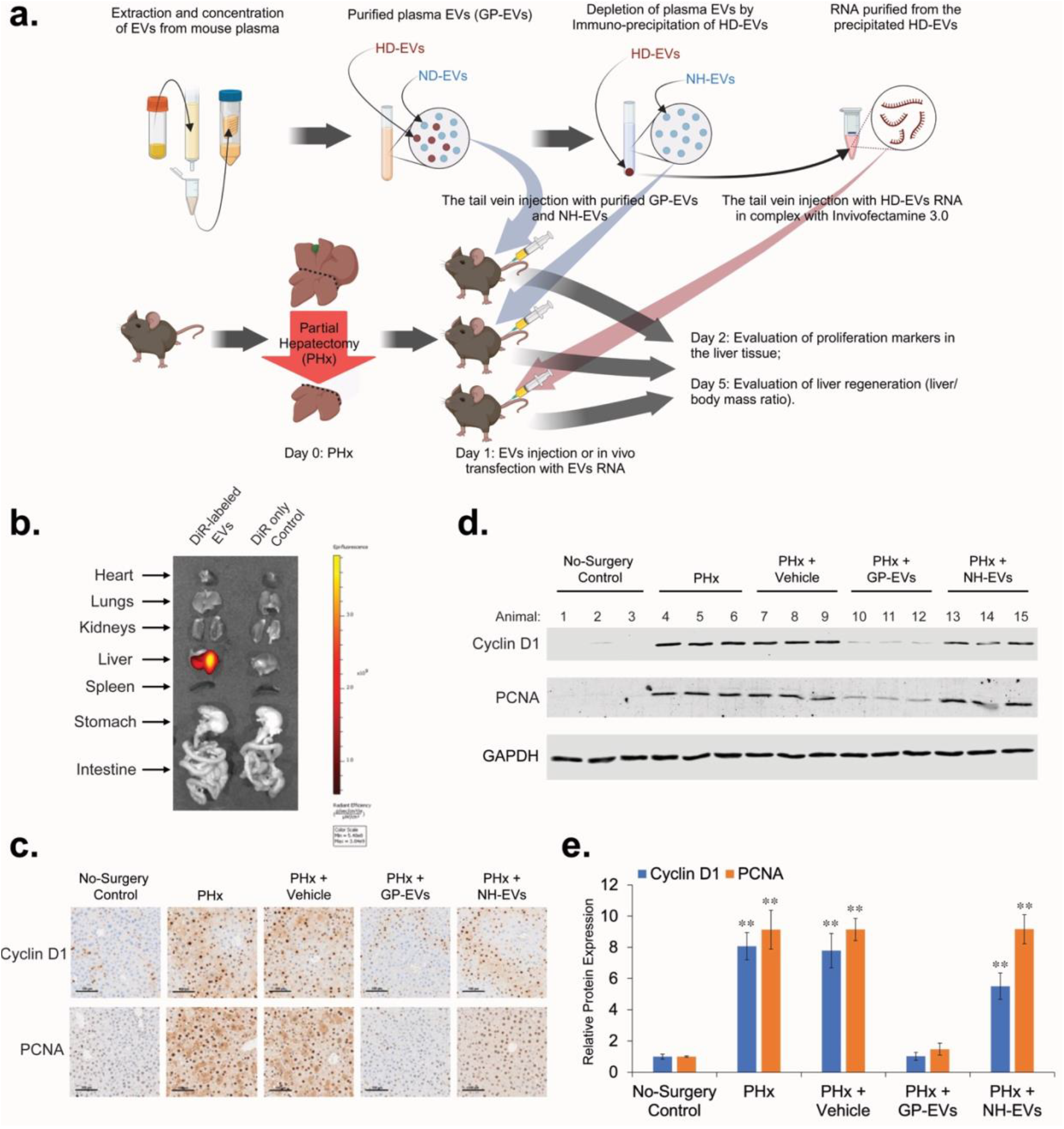
HD-EVs blocks proliferation of mouse hepatocytes in vivo after PHx. **a**. The schematic illustration of the in vivo experiments. **b**. Imaging of the indicated organs for detection of DiR-labeled EVs, 24 hours after IV injection of mice with DiR control only and with DiR-labeled GP-EVs (2×10^11^ particles in 200µL of PBS). **c**. IHC for proliferation markers PCNA and Cyclin D1 in mouse liver 36 hours after PHx. After PHx, different groups of animals received two IV injections (12 hours and 24 hours after PHx) with: i) Vehicle (200µL of PBS) (n = 3); ii) GP-EVs (1×10^12^ particles in 200µL of PBS) (n = 3); iii) ASGR1-neg EVs (1×10^12^ particles in 200µL of PBS) (n = 3). As a positive control, animals didn’t receive any IV injections after PHx (n = 3). As a negative control, intact animals were used (n = 3). Scale bar = 100 µm. **d**. Western blot analysis of liver tissue lysates with Cyclin D1, PCNA, and GAPDH for animal groups showed on Fig. 5c. **e**. Graph with quantification of the Western blot analysis shown in Fig. 5d. n = 3 animals in each group. ** - p-value < 0.001.

For *in vivo* transfection with RNA of the remaining lobes of mouse liver after PHx, we used lipid nanoparticle reagent Invivofectamine™ 3.0. IV injection of Cy™3-labeled Negative Control siRNA in complex with Invivofectamine™ 3.0 demonstrated that 48 hours after injection most of Cy™3-labeled RNA was accumulated in liver tissue (Figure 6a). Some accumulation of Cy™3-labeled RNA was also observed in spleen. IV injection of siRNA targeting mRNA for Factor VII in complex with Invivofectamine 3.0 reagent showed that expression of proteins in liver tissue can be effectively blocked by RNA doses as small as 0.3 mg/kg (Extended data, Figure S2) and this effect can last for several days. To test the effect of RNA from HD-EVs on liver regeneration, we conducted an IV injection of HD-EVs-extracted RNA/Invivofectamine™ 3.0 complex 6 hours after PHx. Forty-eight hours after PHx, IHC and Western blot analyses demonstrated that *in vivo* transfection with HD-EVs-extracted RNA significantly attenuated the PHx-stimulated increase in the expression of proliferation biomarkers Cyclin D1 and PCNA in liver tissue (Figures 6b-d). *In vivo* transfection with the negative control RNA (Scr. RNA), did not affect increase of proliferation markers Cyclin D1 and PCNA after PHx. In additional studies we assessed liver regeneration by calculating the liver/body weight ratio 5 days after PHx. Animals that received IV injection of HD-EVs-extracted RNA/Invivofectamine™ 3.0 complex 6 hours after PHx demonstrated significantly lower liver/body weight ratio compared to animals with no injections after PHx (Figures 6e,f). There was no substantial difference in the liver/body weight ratio at day 5 after PHx between the animals that received an injection of the negative control RNA/Invivofectamine™ 3.0 complex and control animals with no injections after PHx.

**Figure 6.**
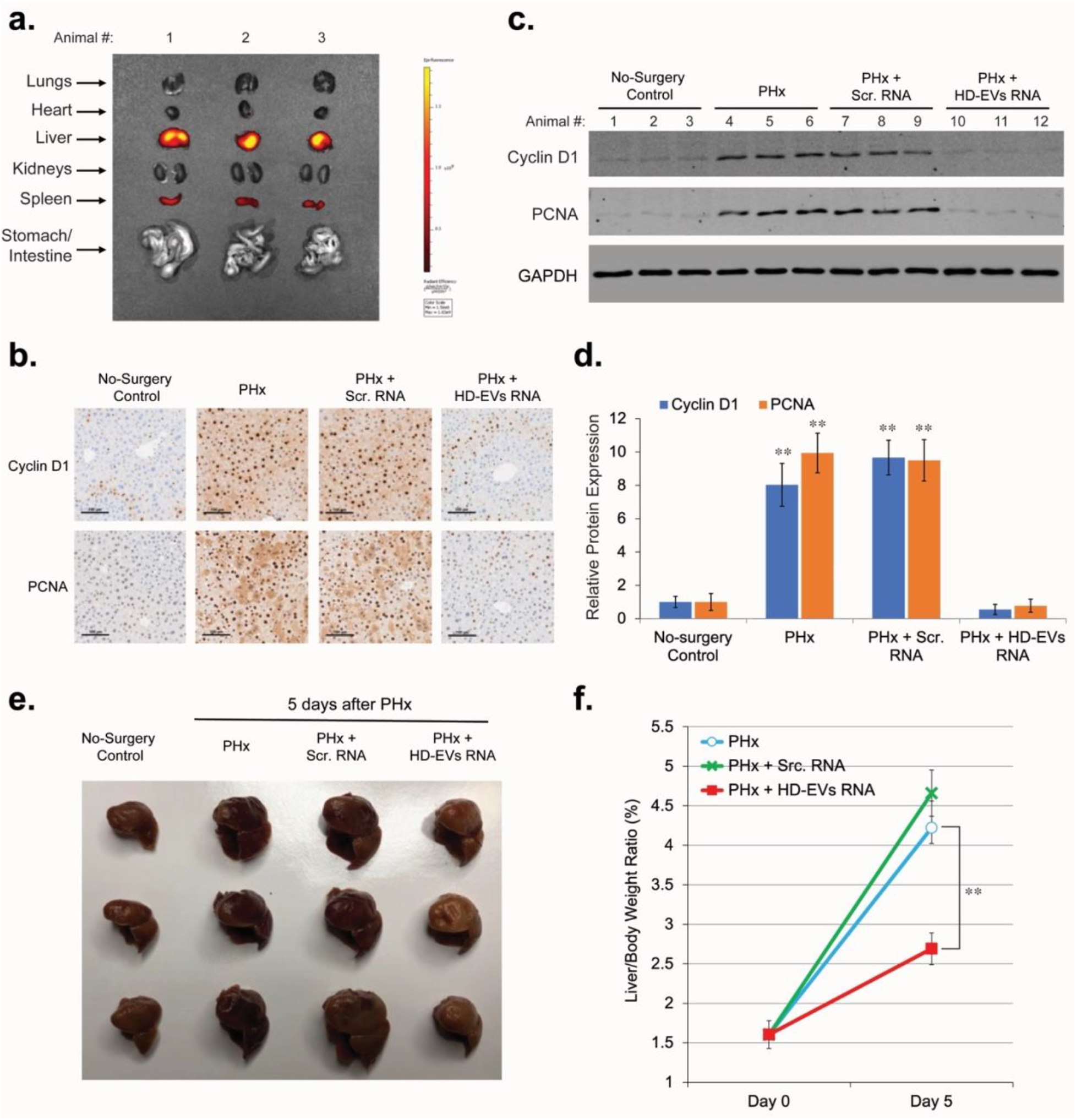
RNA extracted from HD-EVs blocks proliferation of mouse hepatocytes in vivo after PHx. **a**. Imaging of the indicated organs for detection of Cy™3-labeled Negative Control siRNA, 48 hours after IV injection in the complex with Invivofectamine 3.0 Reagent. **b**. IHC for proliferation markers PCNA and Cyclin D1 in mouse liver 48 hours after PHx. 6 hours after PHx, different groups of animals were IV injected with Invivofectamine™ 3.0 Reagent in complex with: i) 1 mg/kg dose of the Negative Control siRNA (n = 3); ii) 1 mg/kg dose of the RNA extracted from HD-EVs (n = 3). As a positive control, animals didn’t receive any IV injections after PHx (n = 3). As a negative control, intact animals were used (n = 3). Scale bar = 100 µm. **c**. Western blot analysis of liver tissue lysates with Cyclin D1, PCNA, and GAPDH for animal groups showed on Fig. 6b. **d**. Graph with quantification of the Western blot analysis shown in Fig. 5c. n = 3 animals in each group. ** - p-value < 0.001. **e**. The median and the right lateral lobes 5 days after PHx from three different groups of animals: i) no additional treatment; ii) animals received IV injection of 1 mg/kg dose of the Negative Control siRNA in complex with Invivofectamine™ 3.0 Reagent, 6 hours after PHx (n = 3); ii) animals received IV injection of 1 mg/kg dose of the RNA extracted from HD-EVs in complex with Invivofectamine™ 3.0 Reagent, 6 hours after PHx (n = 3). The median and the right lateral lobes of the intact animals (no surgery control) are shown as a negative control. **f**. Graph of the quantitative change in the liver/body weight ratio 5 days after surgery for the groups of animals shown in Fig. 6g. n = 3 animals in each group. ** - p-value < 0.001.

## Discussion

Many studies have focused on identification of putative inhibitors of liver cell proliferation (ILCP) and regulation of liver regeneration^31-37^. The detection and characterization of ILCP would enable an understanding of the negative feedback mechanism regulating hepatocyte cell cycle initiation, progression, and completion after a massive liver tissue loss or injury. Although various fractions of plasma or liver extracts have demonstrated a clear inhibitory effect on hepatocyte proliferation, all attempts to isolate and identify the ILCP responsible for this effect have been unsuccessful. Here we show that HD-EVs satisfy most of the previously formulated principal characteristics of ILCP: a) HD-EVs inhibit mitosis of hepatocytes both *in vitro* and *in vivo*; b) action of HD-EVs is dose-dependent, reversible and are not cytotoxic; c) HD-EVs are synthesized by mature liver hepatocytes upon which they act, and circulate in the blood stream; d) HD-EVs are tissue-specific: they are taken up by hepatocytes at a much higher rate than EVs from other sources. We have demonstrated that the decrease in the concentration of HD-EVs in the blood after PHx correlates with the pronounced and coordinated activation of proliferation of almost all hepatocytes in the remaining lobes of the liver. When the liver mass reaches the initial liver/body mass ratio, the plasma HD-EV concentration also reaches the pre-PHx level, which leads to the blocking of hepatocyte proliferation and completion of liver regeneration. Restoration of the initial concentration of HD-EV in the blood by IV injection in the post-PHx period blocks the proliferative activity of hepatocytes and significantly slows down liver regeneration. HD-EVs exhibit the same dose-dependent effect *in vitro*, blocking the proliferation of mouse hepatocytes. It was also shown that RNA isolated from HD-EVs exhibits the same inhibitory effect on hepatocyte proliferation as native HD-EVs. Thus, RNA carried by HD-EVs is an essential component of the mechanism of hepatocyte proliferation inhibition by HD-EVs. Sequence analysis of total RNA extracted from exosomes showed that there was a diverse collection of the RNA species among which miRNAs were the most abundant, making up over 76.20% of all mappable reads compared with only 1.36% of mRNA coding sequences^38^. It is noteworthy that EVs from different cell types carry specific subsets of miRNAs^39^, and miRNAs have sorting sequences that determine their packaging into EVs (EXO-motifs) or retention in cells (CELL-motifs), and that different cell types preferentially use certain sorting sequences, thereby determining the miRNA profile in EVs for a specific cell type^40^. Thus, we suggest that HD-EVs carry a group of miRNAs, which, when returned to hepatocytes, can suppress the activity of the target genes responsible for cell proliferation. Our next goal is the identification of HD-EVs’ miRNAs with ILCP activity and their target genes in hepatocytes.

This study allows us to make another interesting conclusion that the default state of liver hepatocytes is not rest, but proliferation. Constantly ready for proliferation, hepatocytes are held in the G1 phase of the cell cycle by HD-EVs secreted into the blood and recaptured by hepatocytes. When, due to massive loss or injury to liver tissue, the number of functional hepatocytes sharply decreases, the concentration of HD-EVs in the blood drops below a certain threshold, which removes the proliferative block from the remaining hepatocytes and they synchronously enter the cell cycle. Hepatocyte proliferation continues until the initial number of hepatocytes is restored and, as a result, the concentration of HD-EVs released by them in the blood reaches the level of blocking their further proliferation. It is also possible that a similar EVs-based negative feedback mechanism is responsible for the homeostasis of other constantly renewing tissues.

## Methods Summary

### EV isolation from plasma

EVs were extracted from plasma samples by size exclusion chromatography (SEC) method using the qEV columns (Izon Science, Christchurch, New Zealand) as described earlier^30^. Prior to EV isolation, SEC columns were conditioned by washing with freshly filtered (0.1 μm) phosphate-buffered saline (PBS). Thawed plasma was filtered (0.2 μm) added to the sample reservoir and EVs were eluted in PBS, which was added to the sample reservoir as the last of the serum entered the column. For the duration of EV isolation, the volume of PBS in the reservoir was kept below 2 mL. The initial fractions (void volume) of flow-through were discarded, EVs were collected as pooled fractions 1 to 4 (20 mL total) (Figure 2a,b). Resulting pooled EV fractions were mixed by gentle inversion 10 times and concentrated using preconditioned Pierce™ Protein Concentrator PES (100 KDa MWCO, Thermo Fisher Scientific, Waltham, MA, USA) to a final EV concentration 2-3×10^12^ particles/ml of PBS. Samples were then stored at −20°C until thawed, analyzed, or used for cell treatment and injection of animals. After a thaw, plasma EVs were analyzed by nanoparticle tracking analysis, transmission electron microscopy, and Western blot analysis (Figure 2c-e).

### Nanoparticle tracking analysis (NTA)

The analysis was carried out with support of the Microscopy Core of VCU. The concentration and size of exosomes were measured using ZetaView® Nanoparticle Tracking Analysis. All samples were diluted in PBS to a final volume of 2 ml. Ideal measurement concentrations were found by pre-testing the ideal particle per frame value (140–200 particles/frame). The manufacturer’s default software settings for EVs were selected accordingly. For each measurement, three cycles were performed by scanning 11 cell positions each and capturing 80 frames per position under following settings: Focus: autofocus; Camera sensitivity for all samples: 78; Shutter: 100; Scattering Intensity: detected automatically; Cell temperature: 25°C. After capture, the videos were analyzed by the in-built ZetaView Software 8.04.02 SP2 with specific analysis parameters: Maximum area: 1000, Minimum area 5, Minimum brightness: 25. Hardware: embedded laser: 40 mW at 488 nm; camera: CMOS. The number of completed tracks in NTA measurements was always greater than the proposed minimum of 1000 to minimize data skewing based on single large particles.

### Animals, PHx surgery, and EVs injection

The Institutional Animal Care and Use Committee of Virginia Commonwealth University approved all mouse experiments. Male C57BL/6 8-12 weeks old animals were obtained from Charles River Laboratories. Mice housed in animal facilities under specific pathogen-free conditions, receiving humane care according to the criteria of the National Institute of Health Guide for Care and Use of Laboratory Animals. Mice were maintained on a 12-h dark–light cycle and allowed free access to standard food and water. All experiments were conducted during the light cycle. Two-thirds partial hepatectomy (PHx) was described previously^13^. Livers were harvested 36-48 hours after PHx (time-period with the maximum number of hepatocyte mitotic structures), 5 days after PHx, and 10 days after PHx (a time point when the level of the hepatocyte mitotic activity in the regenerated liver is back to normal).

### EV labeling, IVIS, and Flow Cytometry

C57BL/6 mouse plasma (Innovative Research, Novi, MI, USA) was thawed, centrifuged for 30 min at 3000 g and submitted to microfiltration with 0.22-mm filters to remove cell debris and apoptotic bodies. Filtered plasma was then stained for 20 minutes at 37°C in the dark with different lipophilic carbocyanine dyes: 1 μM of DiL dye (1,1’-dioctadecyl-3,3,3’,3’-tetramethylindocarbocyanine, Thermo Fisher Scientific, Waltham, MA, USA), 1 μM DiR dye (1,1’-dioctadecyl-3,3,3’,3’-tetramethylindotricarbocyanine iodide, Thermo Fisher Scientific, Waltham, MA, USA), or 1 μM of DiB dye (CellBrite® Blue Cytoplasmic Membrane Labeling Kit, Biotium, Inc., Fremont, CA, USA). Immediately after staining, EVs were extracted from plasma samples by size exclusion chromatography (SEC) (see sub-chapter “EVs isolation from plasma” for more details), concentrated by using preconditioned Pierce™ Protein Concentrator PES (100 KDa MWCO, Thermo Fisher Scientific, Waltham, MA, USA) to a final EVs concentration 2-3×10^12^ particles/ml of PBS. The collected DiL- or DiR-stained EVs were used fresh for cell culture treatment and cell imaging (DiL-labeled EVs) or for IV injection of animals after PHx (DiR-labeled EVs). Hepatocyte-derived EVs (HD-EVs) were labeled by overnight incubation (rotor, +4C^°^) with Alexa Fluor 647 (AF647)-conjugated anti-mASGR1 Ab (Bioss Antibodies, Woburn, MA, USA) at a dilution 1:500. Flow Cytometry analysis of plasma EVs labeled with DiB and AF647-conjugated anti-mASGR1 Ab was performed using the Cytek® Aurora cytometer (Cytek, Bethesda, MD, USA) and analyzed by SpectroFlo® software. The distribution of EVs labeled with DiR ex vivo was detected by the IVIS® Spectrum system (PerkinElmer, Waltham, USA) as instructed.

### Precipitation of hepatocyte-derived EVs (HD-EVs)

HD-EVs were selectively isolated by combining global plasma EVs (GP-EVs) with Dynabead M280 streptavidin magnetic beads (Thermo Fisher Scientific, Waltham, MA, USA) conjugated with a biotin labeled anti-ASGPR1 polyclonal antibody (Bioss Antibodies, Woburn, MA, USA). Prior to the immunoprecipitation step, the Dynabeads (1 mg) were washed 3 times with 1 mL of filtered (0.1 μm) PBS, separated for 1 minute using the DynaMag-2 magnet, and resuspended in 100 μL of PBS. Washed beads were then incubated with 10 μg of biotinylated anti-ASGPR1 antibody for 1 hour at room temperature with gentle rotation. Antibody-coated beads were separated on the magnet for 2 minutes and washed 4 times with 500 μL of PBS containing 0.1% bovine serum albumin before resuspension in 100 μL of PBS. Isolated plasma EVs (100 μL) were then incubated with the antibody-conjugated beads (100 μL) at 4°C for 24 hours on a rotating mixer. The HD-EVs (ASGPR1+ EVs) bound to Dynabeads were separated out on the magnet for 2 minutes and the depleted supernatant was used for extraction and concentration of non-hepatocyte EVs (NH-EVs) as was described above. As a negative control, EVs precipitation was performed with biotinylated rabbit IgG isotype control (Cell Signaling Technology, Danvers, MA, USA).

### Cell culture, cellular uptake of EVs, and Heparin treatment

Normal mouse hepatocytes (AML-12) were obtained from American Type Culture Collection and were used within 6 months after resuscitation. AML-12 cells were grown in a 1:1 mixture of DMEM/F-12 (Thermo Fisher Scientific, Waltham, MA, USA) supplemented with 10% EVs-depleted FBS (System Biosciences, Palo Alto, CA, USA), 40 ng/ml dexamethasone (Sigma-Aldrich, Inc., St. Louis, MO, USA), and Insulin-transferrin-sodium selenite media supplement (Sigma-Aldrich, Inc., St. Louis, MO, USA). AML-12 cells were propagated in T-75 flasks (VWR, Bridgeport, NJ, USA) and experiments were carried out in 6-well plates (Corning, Glendale, AZ, USA). Twenty-four hours after AML-12 cells were plated, the DiL-labeled EVs were added to the cell media at a final concentration of 2×10^11^ particles/ml. After incubation with labeled EVs for 1-12 hours at +37 °C, cells were washed with cold PBS, fixed with 4% paraformaldehyde for 10 min at +4 °C, and stained with mounting media with DAPI (Ibidi USA, Inc., Fitchburg, WI, USA) for 10 min at +25 °C. To inhibit cellular uptake of EVs, cells and EVs were preincubated with 5 µg/ml Heparin for 4 hours. Cell images were observed by AMG EVOS FL digital inverted microscope (Thermo Fisher Scientific, Waltham, MA, USA).

### Cell cycle analysis

For the cell cycle distribution analysis, AML-12 cells were stained with a FxCycle™ PI/RNase Staining Solution (Thermo Fisher Scientific, Waltham, MA, USA) following the manufacturer’s instructions. Flow cytometry of the stained cells was performed with a FACSCanto II flow cytometer (BD Biosciences, Franklin Lakes, NJ, USA).

### Immunohistochemistry (IHC)

The liver samples were fixed in 10% Neutral Buffered Formalin (Sigma-Aldrich, Inc., St. Louis, MO, USA) at +4 °C. Then the fixed materials were dehydrated with 70% ethanol, embedded with paraffin, and sectioned at 5 μm for staining. The IHC staining and analysis was carried out by the Tissue and Data Acquisition and Analysis Core of VCU Massey Comprehensive Cancer Center.

### Western blot analysis

For protein analysis all samples were loaded and separated by SDS-PAGE gel and transferred to nitrocellulose membranes. The membranes were exposed to antibodies at specific dilutions. Primary antibodies used for WB: anti-CD63 (dilution 1:1000, Thermo Fisher Scientific, Waltham, MA, USA), anti-TSG101 (dilution 1:500, Cell Signaling Technology, Danvers, MA, USA), anti-Albumin (dilution 1:1000, Cell Signaling Technology), anti-Cyclin D1 (dilution 1:1000, Cell Signaling Technology), anti-GAPDH (dilution 1:2000, Cell Signaling Technology), anti-PCNA (dilution 1:1000, Cell Signaling Technology). Specific protein bands were detected using infrared-emitting conjugated secondary antibodies: anti-rabbit DyLight™ 800 4X PEG Conjugate (dilution 1:10,000, Cell Signaling Technology). WB images were generated and analyzed using the Odyssey CLx imaging system and iS Image Studio 5.2.5 software (Li-Cor).

### Transmission Electron Microscopy (TEM) of isolated EVs

Isolated EVs were fixed and prepared for the TEM as previously described^41,42^. Briefly, purified EVs resuspended in PBS were fixed with an equal volume of 4% Paraformaldehyde (PFA) (Thermo Fisher Scientific, Waltham, MA, USA) containing 0.1M sodium cacodylate buffer (Avantor, Allentown, PA, USA). The final concentration of this solution is 2% PFA, and samples were kept in fixative for up to 48 hours at +4°C until further processed. Eight μl drops of each fixed sample were deposited onto a Formvar-coated grid (composed of copper with carbon) (Sigma-Aldrich, Inc., St. Louis, MO, USA) to allow the vesicles to attach to the grid for 30 minutes at RT. The grids were then washed with PBS and transferred onto a drop of 1% glutaraldehyde (Sigma-Aldrich, Inc., St. Louis, MO, USA) in 0.1M sodium cacodylate buffer for 5 minutes. The grids were then washed with diH_2_O ×5 times over the course of 2 minutes. After washing, grids were transferred onto a drop of fresh 0.5% aqueous uranyl acetate solution (Avantor, Allentown, PA, USA) for 3 minutes for staining. Grids were then blotted to remove excess liquid, air-dried (typically overnight), and stored in the dark until imaged. Following sample preparation, EVs were imaged on the JEOL JEM-1400 Plus TEM (JEOL USA, Inc., Peabody, MA, USA) with sCMOS OneView camera (Gatan, Inc., Pleasanton, CA, USA) at 100kV. The TEM analysis was carried out support of the Microscopy Core of VCU.

### RNA extraction and estimation

MicroRNA was purified from neuron derived EV samples using the MiRNeasy Micro Kit (Qiagen) according to the manufacturer’s protocol with a few modifications to maximize the recovery of small RNA from EVs. Extracted EVs (in 150 μL of PBS) were homogenized with 750 μL of TRIzol™ LS reagent (Invitrogen), followed by 200 μL of chloroform. Each sample was vortexed for 60 s and incubated at room temperature for 5 min. Phase separation was performed by centrifugation at 12,000×*g* for 15 min at 4 °C. Three hundred µL of the upper aqueous phase were transferred to a new tube. To stimulate RNA precipitation, glycogen (5 mg/mL) (Invitrogen) was added to the aqueous phase before being mixed with 750 μL of cold 100% molecular grade Ethanol. Tubes were vortexed for 30 s and incubated at − 20 °C for 1 hour. After precipitation step, samples were transferred to a Qiagen RNeasy® Mini spin column in a collection tube followed by centrifugation at 15,000 × g for 30 s at room temperature. Then, the Qiagen RNeasy® Mini spin columns were washed three times according to the standard protocol. After the last wash with 70% Ethanol, the Qiagen RNeasy® Mini spin columns were left uncapped for 5 min to allow the column to dry. Next, 20 μL of RNase-free pre-warmed (+65°C) water was added to the dry columns, and after 5 min of incubation, total EVs RNA was eluted in a new RNase free collecting tubes by 1 min centrifugation at 15,000 × g. To increase the amount of eluted RNA, columns were reloaded with the eluent and centrifuged again. The eluents with total EVs RNA were stored at −80 ° C. The concentration and purity of the EVs RNA was measured using NanoDrop ND-1000 spectrophotometer (Thermo Fisher Scientific, Waltham, MA, USA).

### RNA transfection *in vitro* and *in vivo*

For in vitro transfection, mouse hepatocytes (AML-12 cell line) were distributed onto 6-well plates at a concentration of 2 × 10^5^ cells/well and incubated overnight to allow cell adherence. 24 hours after plating, RNA transfection was performed in the presence of Lipofectamine RNAiMAX (Invitrogen) according to manufacturer’s protocol. As a negative control, cells were transfected with *Silencer*™ Negative Control No. 1 siRNA (Thermo Fisher Scientific, Waltham, MA, USA). Two days after transfection, cells were trypsinized, washed, and prepared for Western blot and Cell Cycle analysis. For *in vivo* transfection, RNA/ Invivofectamine™ 3.0 Reagent (Thermo Fisher Scientific, Waltham, MA, USA) complex was prepared according to the manufacturer’s protocol. Six hours after PHx, animals received tail vein injection with 200 μL volume of RNA/Invivofectamine™ 3.0 Reagent complex. For visualization of RNA distribution in animals’ organs, and as a negative control, *Silencer*™ Cy™3-labeled Negative Control No. 1 siRNA was used (Thermo Fisher Scientific, Waltham, MA, USA).

### Statistical significance

Statistical significance was determined using a 2-sided Student’s t-test (Fig. 1c,f; 6f) or Analysis of Variance (ANOVA) (Fig. 3g; 4d; 5e; 6d). P values less than 0.05 were considered significant.

## Acknowledgments

TEM and Nanoparticle Tracking Analysis were performed at the VCU Microscopy Facility. Flow Cytometry of the labeled EVs was performed at the VCU Flow Cytometry Shared Resource. IVIS analysis was performed at VCU Cancer Mouse Models Core Laboratory. IHC was performed by Tissue and Data Acquisition and Analysis Core of VCU Massey Comprehensive Cancer Center. All these services are supported, in part, by funding from NIH-NCI Cancer Center Support Grant P30 CA016059. Dr. Yakovlev and Dr. Rabender were supported by the internal fund of the VCU Massey Cancer Center.

## Author contributions

Conceptualization and design of the study: VY. VY and CR performed a mouse surgery, animal post-surgery care, collection of the plasma and liver samples. VY and MM performed cell culture experiments. VY and MM performed bio-molecular analyzes. Data analysis: VY, RM. Writing the original draft: VY; all other authors commented on and refined the manuscript. Data visualization: VY. All authors have carefully read the paper and approved the final manuscript.

