## Extended data, Figure S1 for "Hepatocyte-derived extracellular vesicles regulate liver regeneration after partial hepatectomy"

**Extended data, Figure S1: Inhibition of EVs uptake by Heparin.** AML-12 cells and EVs were incubated with different concentrations of Heparin for 2 hours. At the end of the incubation time, the cells were washed, fixed, and stained with mounting media with DAPI. Scale bar = 200 µm.

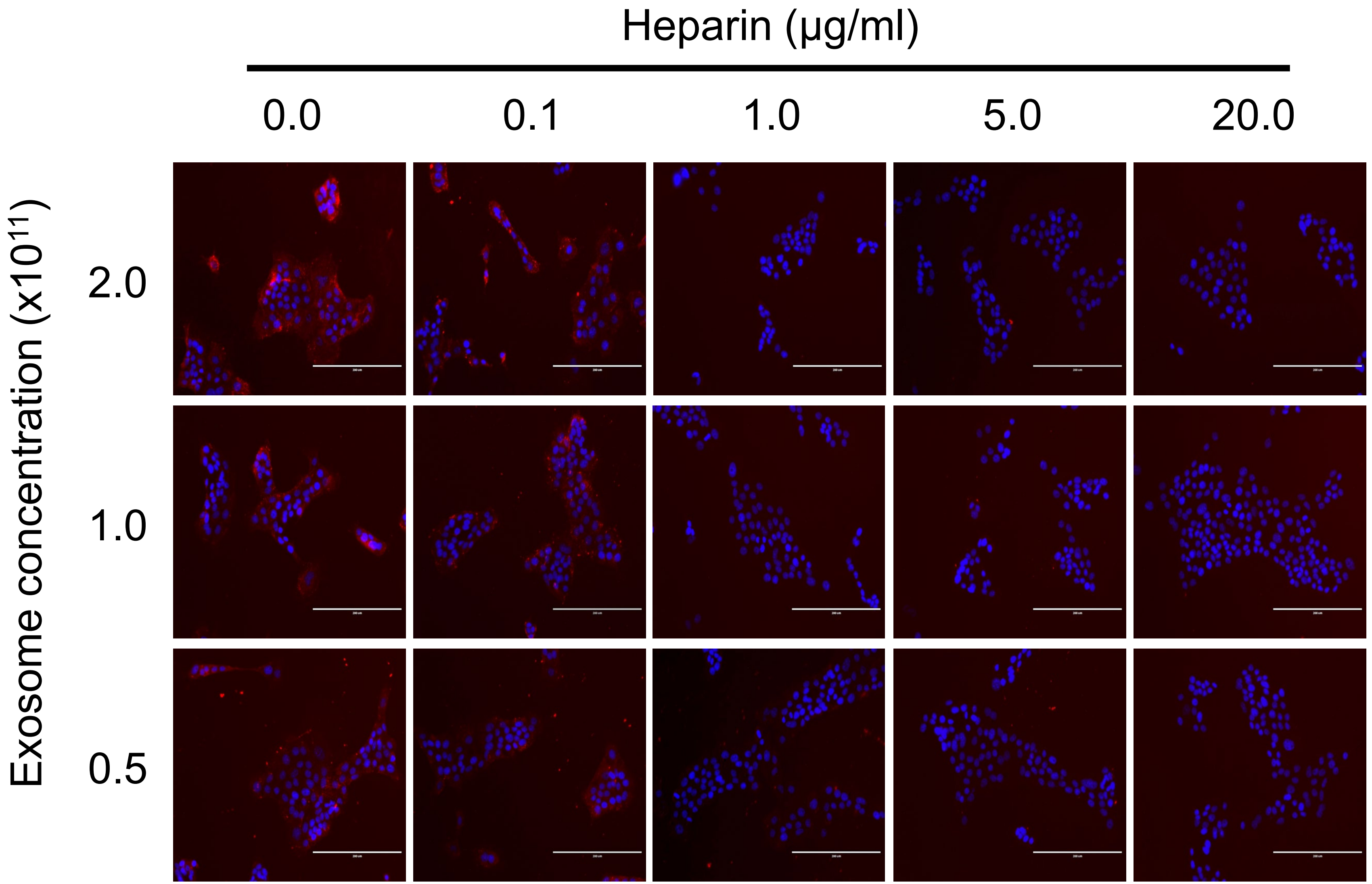
