## Extended data, Figure S2 for "Hepatocyte-derived extracellular vesicles regulate liver regeneration after partial hepatectomy"

**Extended data, Figure S2: In Vivo inhibition of Factor VII protein expression in the mouse liver.** Animals were injected via tail vein (IV) with 0.3-0.5 mg/kg dose of siRNA Factor VII in complex with Invivofectamine 3.0 Reagent. Control animals were injected with 0.5 mg/kg dose of Negative Control siRNA. Factor VII protein level in liver tissue was estimated 24 and 48 hours after injections by Western Blot. Western Blot with GAPDH was used as normalization control.

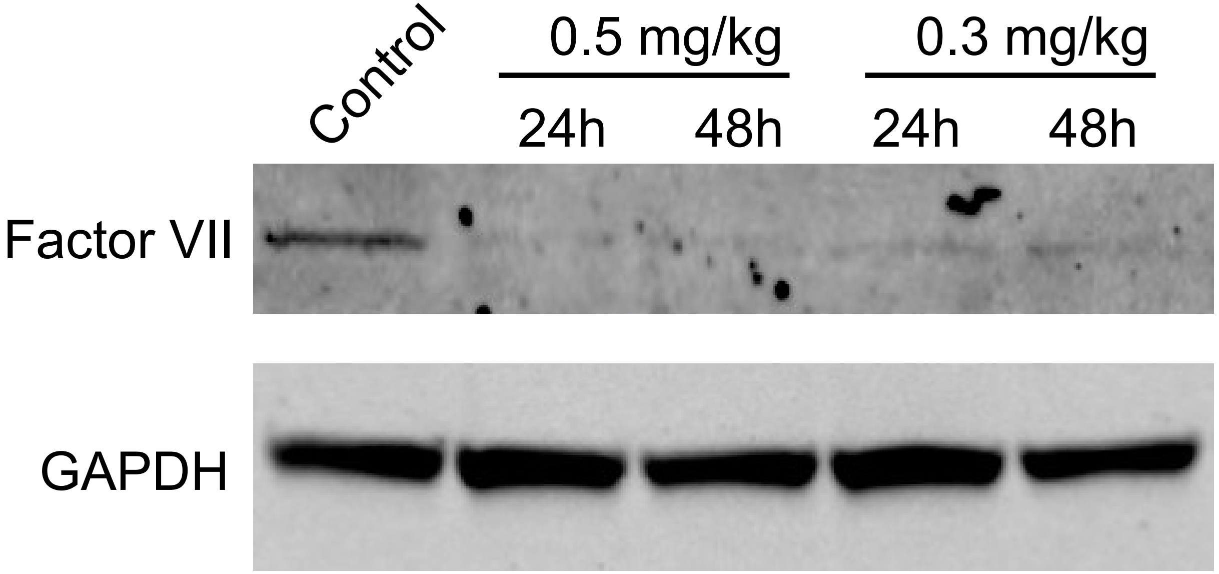
